# Intrinsically disordered regions contribute promiscuous interactions to RNP granule assembly

**DOI:** 10.1101/147561

**Authors:** David S. W. Protter, Bhalchandra S. Rao, Briana Van Treeck, Yuan Lin, Laura Mizoue, Michael K. Rosen, Roy Parker

## Abstract

Eukaryotic cells contain large RNA-protein assemblies referred to as RNP granules, whose assembly is promoted by both traditional protein interactions and intrinsically disordered protein domains. Using RNP granules as an example, we provide evidence for an assembly mechanism of large cellular structures wherein specific protein-protein or protein-RNA interactions act together with promiscuous interactions of intrinsically disordered regions (IDRs). This synergistic assembly mechanism illuminates RNP granule assembly, and explains why many components of RNP granules, and other large dynamic assemblies, contain IDRs linked to specific protein-protein or protein-RNA interaction modules. We suggest assemblies based on combinations of specific interactions and promiscuous IDRs are common features of eukaryotic cells.

## INTRODUCTION

Eukaryotic cells contain a variety of non-membrane bound RNA-protein assemblies, collectively referred to as RNP granules. Such RNP granules include the nucleolus and Cajal body in the nucleus, as well as stress granules and P-bodies in the cytosol (Spector, 2006). RNP granules are generally highly dynamic, as judged by FRAP of their protein components, and exhibit liquid-like behaviors, such as flowing, fusing, and rapid reorganization of internal components (Brangwynne, 2013; Brangwynne et al., 2009). RNP granules are thought to assemble through a process referred to as liquid-liquid phase separation (LLPS) wherein RNA molecules provide binding sites for RNA binding proteins that interact with themselves or other RNA binding proteins to create a larger multivalent assembly (Elbaum-Garfinkle et al., 2015; Feric et al., 2016; Kaiser et al., 2008; Mitrea et al., 2016; Nott et al., 2015; Pak et al., 2016; Patel et al., 2015; Riback et al., 2017; Weber and Brangwynne, 2012; Zhang et al., 2015). Some of the interactions that drive RNP granule assembly are well defined interactions between folded proteins, or folded protein domains and short linear motifs (SLiMs) (Decker et al., 2007; Jonas and Izaurralde, 2013; Kedersha et al., 2016; Ling et al., 2008; Mitrea et al., 2016; Tourriere, 2003). Since these interactions require folded protein structures and/or extended linear motifs that interact in a stereospecific manner, we refer to these interactions as specific interactions.

The IDRs of RNA binding proteins have been highlighted as drivers of RNP granule assembly for three reasons. First, genetics indicate that IDRs can be important for assembly of RNP granules or localization of granule components (Decker et al., 2007; Feric et al., 2016; Gilks et al., 2004; Hennig et al., 2015; Kato et al., 2012). Second, RNP granules are often enriched in proteins with IDRs (Decker et al., 2007; Jain et al., 2016; Kato et al., 2012; Kedersha et al., 2013; Reijns et al., 2008). Finally, IDRs are often (but not always) both necessary and/or sufficient for LLPS of granule proteins *in vitro*, forming structures that resemble RNP granules *in vivo -* (Elbaum-Garfinkle et al., 2015; Lin et al., 2015; Molliex et al., 2015; Nott et al., 2015; Patel et al., 2015; Smith et al., 2016; Zhang et al., 2015).

An unresolved issue is how IDRs contribute to RNP granule assembly, and how IDR based assembly mechanisms integrate with specific protein-protein and protein-RNA interactions to promote RNP granule formation. The literature suggests three non-mutually exclusive models by which IDRs could contribute to LLPS *in vitro* and RNP granule formation *in vivo*. First, some experiments *in vitro* suggest that IDRs promote LLPS via weak binding, utilizing electrostatic, cation-π, dipole-dipole and π-π stacking interactions (Brangwynne et al., 2015; Lin et al., 2016; Nott et al., 2015; Pak et al., 2016). Charge patterning also appears to play an important role, wherein like-charged amino acids are clustered together within an IDR. Scrambling these charges across the length of an IDR has been observed to impair LLPS both *in vitro* and *in vivo* (Nott et al., 2015; Pak et al., 2016). Because these interactions only require a few amino acids, and do not require any stereospecific arrangement, they would be anticipated to occur between an IDR and many other proteins, including other IDRs. Indeed, charge patterning specifically has been proposed to mediate interactions between IDRs and cellular proteins (Pak et al., 2016). For this reason, we refer to the above types of IDR interactions as nonspecific. These interactions will also be promiscuous, because they will be relatively indiscriminate with respect to binding partners. A second possibility is that elements within some IDRs interact in a specific manner involving local regions of secondary structure. For example, there is a correlation with how mutations in hnRNPA2 affect binding of its C-terminal disordered domain to beta-strand rich hydrogels, and the recruitment of those hnRNPA2 domain variants to LLPS of wild-type hnRNPA2 (Xiang et al., 2015). Similarly, a locally formed α-helix in TDP-43 can mediate LLPS through homotypic interactions (Conicella et al., 2016). Finally, it is likely that a subset of IDRs are also promiscuous RNA binding proteins since they can be rich in positive charges, some IDRs can cross link to mRNA *in vivo*, and some IDRs can bind RNA *in vitro* (Lin et al., 2015; Lyons et al., 2014; Mayeda et al., 1994; Molliex et al., 2015).

Given the promiscuous nature of IDR interactions, we hypothesized that such IDR based interactions alone would be susceptible to other highly abundant proteins in cells, and therefore insufficient to drive LLPS and the assembly of an RNP granule *in vivo*. In the context of the protein-rich cellular environment other proteins would compete for binding to the IDRs and thereby prevent their forming a defined assembly. Moreover, even the ability of some IDRs to form specific local structure based interactions might be impaired by competition with other proteins in the cell. Instead, to account for the contributions from both IDRs and specific interactions to RNP granule assembly, we hypothesized that IDRs would reinforce assemblies that contained specific assembly interactions. Effectively, specific interactions would concentrate the IDRs and strengthen their interactions through additive binding energies (Jencks, 1981), either biasing their promiscuous interactions toward components of the assembly, or promoting the formation of specific interactions between the IDRs. In this way, IDR-based interactions could contribute to the energetics of assembly.

Here we provide several observations that RNP granule assembly gains selectivity from specific protein-protein and protein-RNA interactions, and that promiscuous binding of IDRs to proteins and possibly RNA enhances these assemblies. First, we observe that LLPS driven by IDRs *in vitro* is inhibited by other proteins. Second, in cells we observe that IDRs of granule components are often neither required nor sufficient to target proteins to RNP granules. Third, we demonstrate that *in vitro* LLPS driven by specific protein-RNA interactions is enhanced by adding promiscuously interacting IDRs, and the assembly of yeast P-bodies in cells is promoted by nonspecific IDRs in conjunction with specific interactions. Thus, RNP granules assemble primarily by specific interactions, which can be enhanced by IDRs capable of either promiscuous, or weak specific interactions based on small structural elements that become effective at high local concentrations. We suggest that this general assembly mechanism may be shared by other macromolecular complexes rich in IDRs.

## RESULTS

### Several proteins inhibit LLPS driven by IDRs *in vitro*

We hypothesized that IDRs of RNA binding proteins might not be sufficient to drive LLPS in the presence of other proteins similar to the intracellular environment, despite the observation that such IDRs are capable of undergoing LLPS as purified proteins (Elbaum-Garfinkle et al., 2015; Lin et al., 2015; Molliex et al., 2015; Nott et al., 2015; Patel et al., 2015; Smith et al., 2016; Zhang et al., 2015). Our hypothesis was based on the observations that LLPS driven by IDRs *in vitro* are thought to occur by weak electrostatic, dipolar interactions as well as interactions involving aromatic groups (reviewed in Brangwynne et al., 2015). Since these interactions are nonspecific, they are likely relatively promiscuous and could, in principle, occur between an IDR and other IDRs or with many other proteins. Moreover, even IDRs that have homotypic interactions based on local structural elements might be sensitive to other proteins and be most efficient at forming such specific assemblies only when concentrated by specific interactions (Xiang et al., 2015; Conicella et al., 2016). Thus, we asked whether IDR driven LLPS *in vitro* would be inhibited in the presence of other polypeptides, which would be analogous to the interior of the cell.

To test whether an IDR can promote LLPS in the presence of other proteins, we induced LLPS of either full-length hnRNPA1_Δhexa_ or only the hnRNPA1_IDR_ region (amino acids 186 to 300, **Figure 1A**) by dilution into lower salt (37 mM NaCl) (Lin et al., 2015) in the presence of increasing amounts of bovine serum albumin (BSA). We used the Δhexa-peptide variant of the full-length hnRNPA1 protein as it is less prone to forming amyloid fibers during purification and analysis, and behaves similarly to the wild-type protein with regards to LLPS (Lin et al., 2015; Molliex et al., 2015). The fluorescently conjugatable SNAP tag was fused to both proteins to visualize droplets.

**Figure 1.**
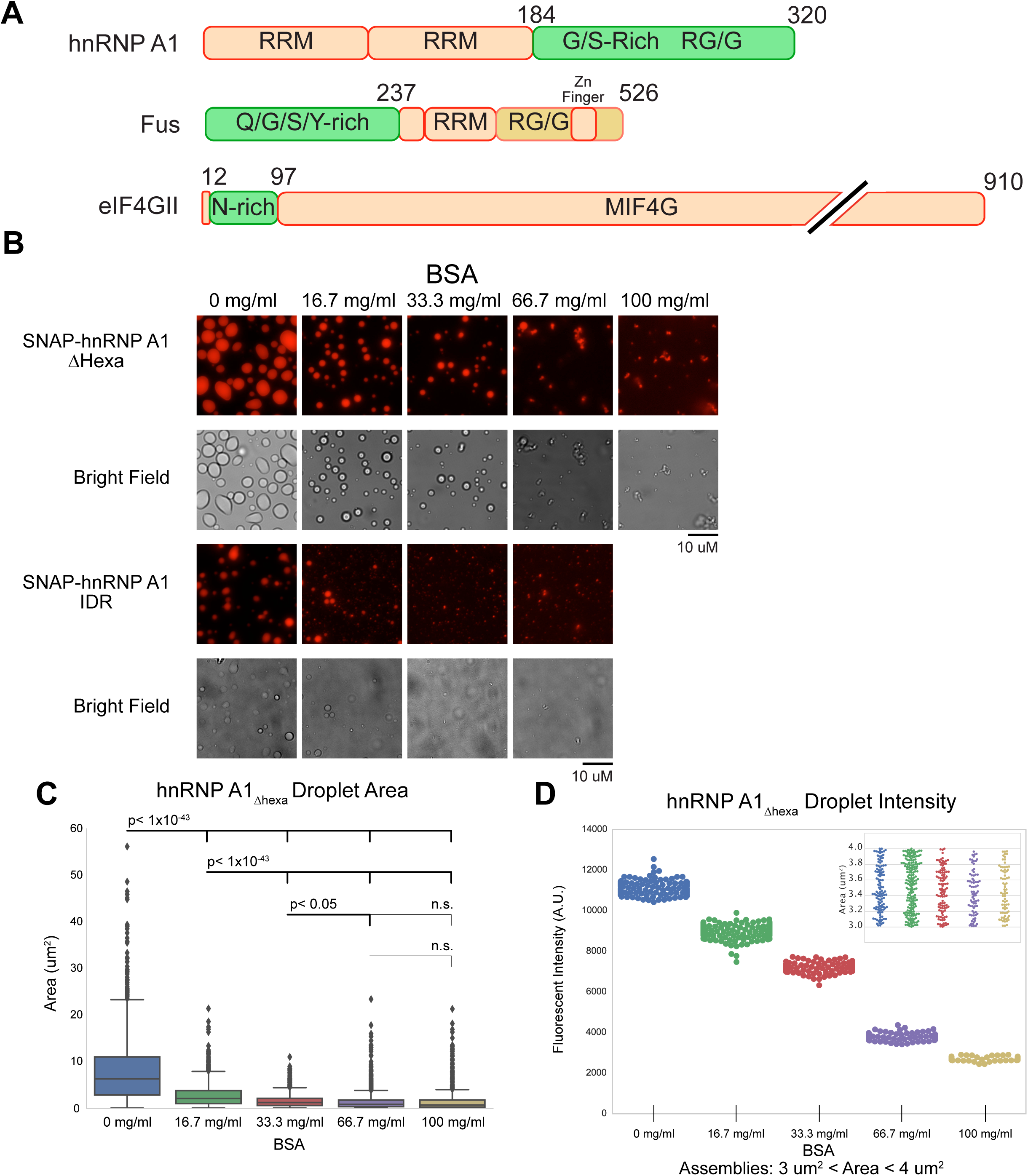
Competitor Proteins Disrupt IDR-Driven Phase Separations. (A) Domain structure of hnRNP A1, FUS, and eIF4GII (B) Fluorescent and bright-field microscopy images of phase separated droplets formed at 37.5 mM NaCl by SNAP-hnRNP A1_Δhexa_ and SNAP-hnRNP A1_IDR_ with the indicated concentrations of BSA. Images are each independently scaled. (C) Quantification of structure size for hnRNP A1_Δhexa_ from (B), significance calculated with Welch’s *t*-test for unequal size and variance. (D) Quantification of the intensity of all structures between areas of 3 um and 4 um for hnRNP A1_Δhexa_ from (B). These subsets of droplets have roughly equal distributions of size (inset)

As the concentration of BSA increased, LLPS for both full-length SNAP-hnRNPA1_Δhexa_ and the SNAP-hnRNPA1_IDR_ was inhibited (**Figure 1B**). At higher BSA concentrations, we observed the formation of aggregated hnRNPA1_Δhexa_ and hnRNPA1_IDR_ that contrast with the liquid droplets seen in the absence of BSA (**Figure 1B**). As BSA concentrations increase, droplet sizes decrease and no large droplets form (**Figure 1C**). Interestingly, by looking at a subset of droplets of similar size across all BSA concentrations, we noticed that as BSA concentrations increased the intensities of hnRNPA1_Δhexa_ droplets decreased (**Figure 1D**). The distribution of areas was approximately equal between samples (**Figure 1D inset**). The partition coefficient of LLPS (the ratio of protein within the concentrated phase versus within the dilute phase) is a measure of the equilibrium between the two states. Therefore, we interpret this decrease in intensity to mean that BSA shifts the phase separation equilibrium such that it is less favorable for hnRNP A1_Δhexa_ to exist within the concentrated phase. At higher BSA concentrations the equilibrium shifts such that hnRNPA1_Δhexa_ is below the critical concentration for LLPS. Thus, BSA is an inhibitor of LLPS driven by hnRNPA1 or its IDR alone under these conditions.

To determine if this inhibitory effect is unique to hnRNPA1 and BSA, we examined how BSA, lysozyme, and RNase A affected LLPS driven by the IDRs of hnRNPA1, FUS, or eIF4GII, all of which have been reported to undergo LLPS at low salt or low temperature (Lin et al., 2015; Molliex et al., 2015; Patel et al, 2015). We observed that LLPS of FUS_IDR_ (amino acids 1-237), eIF4GII_IDR_ (amino acids 13-97), or the hnRNPA1_IDR_ (184-320) (**Figure 1A)** were also inhibited by the presence of BSA, lysozyme or RNase A **(Figure 2A)**.

**Figure 2.**
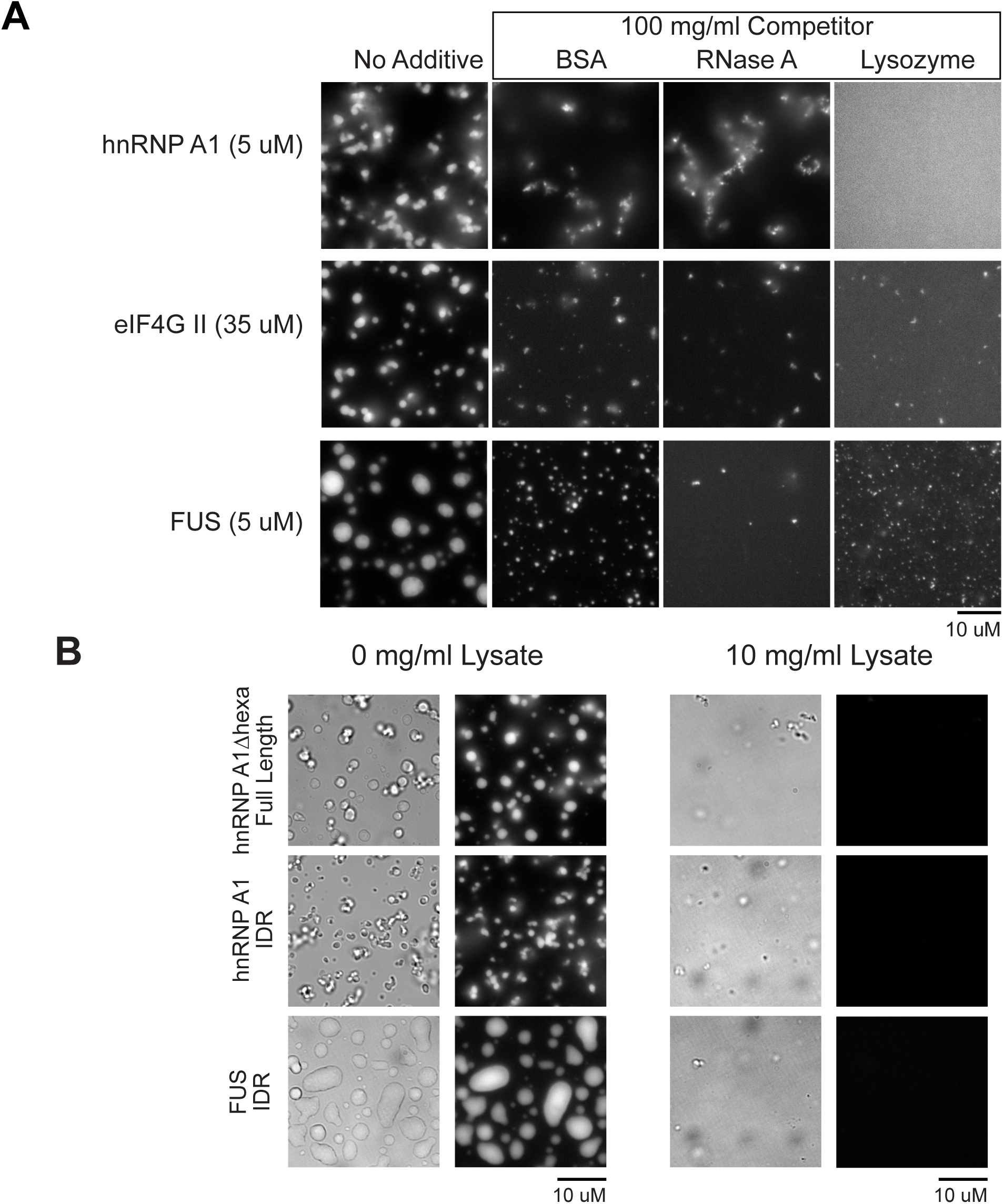
Globular Proteins Are Recruited to IDR-driven LLPS Droplets. (A) Fluorescent microscopy images of phase separated droplets formed at 37.5 mM NaCl by hnRNP A1_IDR_, SNAP-FUS_IDR_, and SNAP-eIF4GII_IDR_ in the absence or presence of 100 mg/ml BSA, lysozyme, and RNase A. (B) Fluorescent microscopy images of phase separated droplets formed at 37.5 mM NaCl by SNAP-hnRNP A1_Δhexa_, hnRNP A1_IDR_, and SNAP-FUS_IDR_, in the absence or presence of approximately 10 mg/ml yeast lysate.

To more closely mimic the cellular environment, we examined whether IDRs or IDR containing proteins could undergo LLPS in the presence of yeast lysate, which had been previously depleted of small metabolites and exchanged into droplet-forming buffer via desalting columns. We observed that LLPS of hnRNPA1_Δhexa_, hnRNPA1_IDR_, and FUS IDR are all strongly impaired in yeast lysates, which contained approximately 10 mg/ml protein (**Figure 2B**). Yeast lysates are our closest approximation of the cellular environment, and we find that even lysates 1/10^th^ as concentrated as the cell (Milo, 2013) strongly impair LLPS of IDRs. Thus, phase separation of multiple IDRs is sensitive to competition from other molecules within the cell.

### Competitor proteins inhibit LLPS *in vitro* by interacting with IDRs

What is the mechanism by which competitor proteins inhibit IDR-driven LLPS *in vitro*? One possibility is that BSA, lysozyme, and RNase A share some specific property or structural feature that inhibits LLPS of these IDRs. This is unlikely as BSA, lysozyme, and RNase A are structurally unrelated, and vary in size (66.4, 14.3, and 13.7 kDa respectively) and pI (5.3, 11.35, and 9.6, respectively). A second possibility is that any crowding agent will inhibit LLPS under these conditions. However, we observe that LLPS driven by hnRNPA1_IDR_ is stimulated by the crowding agents Ficoll and PEG, with phase separation occurring at higher ionic strengths and lower protein concentrations than without crowding agents **(Figure S1A)** (see also Lin et al., 2015; Molliex et al., 2015).

A third possibility is that these competitor proteins compete for promiscuous interactions between IDRs and thereby disrupt LLPS. A prediction of this model is that at low concentrations, insufficient to block LLPS, the competitor proteins would be recruited into the phase separated droplets (due to interactions with the IDR). To test this possibility, we examined the recruitment of fluorescent BSA or lysozyme into droplets formed by IDRs. At low concentrations both proteins were recruited to IDR-driven droplets without disrupting the assemblies. For example, at 500 nM concentration FITC-BSA was strongly enriched in droplets of hnRNPA1_Δhexa_ **(Figure 3A)**. FITC-Lysozyme was also recruited (**Figure 3A**). Similarly, droplets of eIF4GII_IDR_, hnRNPA1_IDR_, and FUSIDR all recruited both FITC-BSA and FITC-lysozyme **(Figure 3B).** This suggests that these IDRs can interact with both BSA and lysozyme, consistent with the idea that competitor proteins could compete with the weak interactions that mediate LLPS.

**Figure 3.**
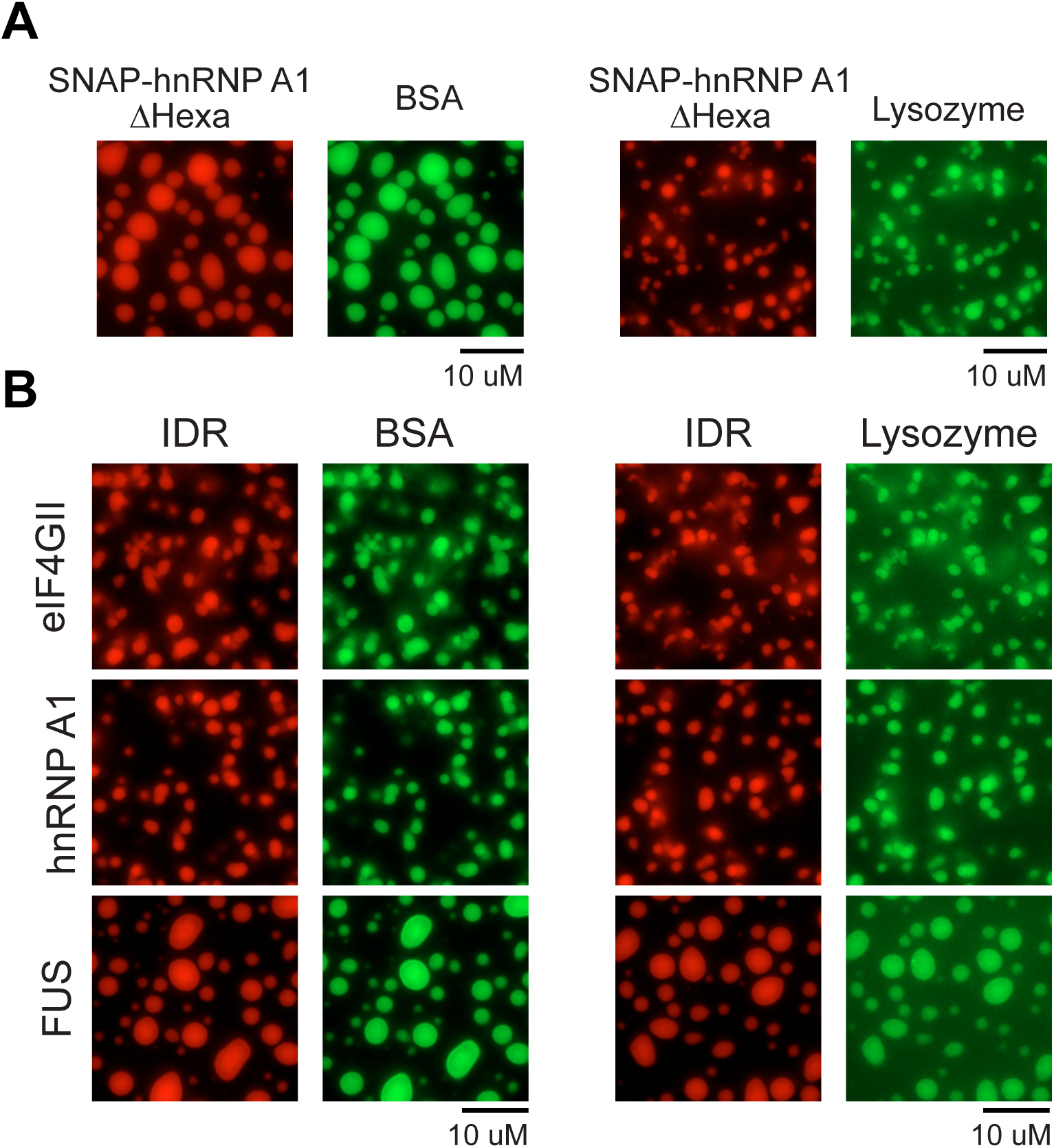
Globular Proteins Are Recruited to IDR-driven LLPS Droplets. (C) Fluorescence microscopy images of phase separated droplets formed at 37.5 mM NaCl by 25 uM SNAP-hnRNP A1_Δhexa_ (red) and 500 nM FITC-labeled BSA (green) or FITC-labeled Lysozyme. (D) Fluorescence microscopy images of phase separated droplets formed by SNAP-eIF4GII_IDR_ (35 uM), SNAP-hnRNP A1_IDR_ (5.25 uM), or SNAP-FUS_IDR_ (5 uM) in the presence of either 10 nM FITC-BSA or 100 nM FITC-Lysozyme.

The above evidence suggests that competitor proteins can interact with IDRs, both because these proteins are recruited into phase-separated droplets and because they inhibit LLPS at higher concentrations. Since these proteins were chosen at random and have diverse physical properties, and LLPS is also inhibited by metabolite-depleted cell lysates, we suggest that IDRs by themselves are likely to be susceptible to such nonspecific interactions in the more complex cellular environment. Therefore, in many cases, promiscuous interactions of IDRs are unlikely to be sufficient for RNP granule assembly in cells.

### IDRs can enhance LLPS driven by specific interactions in the presence of competitor proteins

The *in vitro* results above suggest that IDR-IDR interactions are susceptible to competition by the complex protein mixture in the cell. However, IDRs are enriched in RNP granule proteins (Decker et al., 2007; Jain et al., 2016; Kato et al., 2012; King et al., 2012; Reijns et al., 2008), and IDRs can play a role in RNP granule assembly (e.g. Decker et al., 2007; Gilks et al., 2004; Wang et al., 2014). In some cases, IDRs contain SLiMs that are important for assembly of RNP granules (reviewed in Jonas and Izaurralde, 2013). However, since there are cases wherein one IDR can functionally substitute for another in RNP granule assembly (Decker et al., 2007; Gilks et al., 2004), a more generic role for IDRs in RNP granule assembly is also likely.

We hypothesized that IDRs in proteins that also make specific interactions could provide promiscuous, nonspecific interactions that stabilize an RNP granule by acting together with the specific interactions. By concentrating the IDRs through specific interactions, promiscuous IDR-based interactions are biased to other components of the assembly. In this model, specific interactions and nonspecific interactions both donate binding energy that promotes LLPS. This model makes two predictions that we first tested *in vitro*.

First, the model predicts that LLPSs driven by specific interactions should be less susceptible to the interference from other competitor proteins, and may even be enhanced, given that high concentrations of such proteins can serve as crowding agents. Consistent with this view, we have shown that the LLPS driven by the specific interaction of an RNA binding protein, poly-pyrimidine tract binding protein (PTB), with RNA is promoted by BSA (Lin et al., 2015), an observation reproduced here. For example, while SNAP-tagged PTB and RNA showed limited assembly when mixed together at concentrations of 20 uM and 1.6 uM, respectively, the addition of 100 mg/ml BSA induced robust phase separation at these concentrations **(Figure 4A)**. Consistent with this effect being due to molecular crowding, the PTB-RNA LLPS is also stimulated by PEG or Ficoll, two additional crowding agents **(Figure 4A).** Thus, the specific PTB-RNA interactions are not outcompeted by BSA, allowing the crowding effect of BSA to dominate.

**Figure 4.**
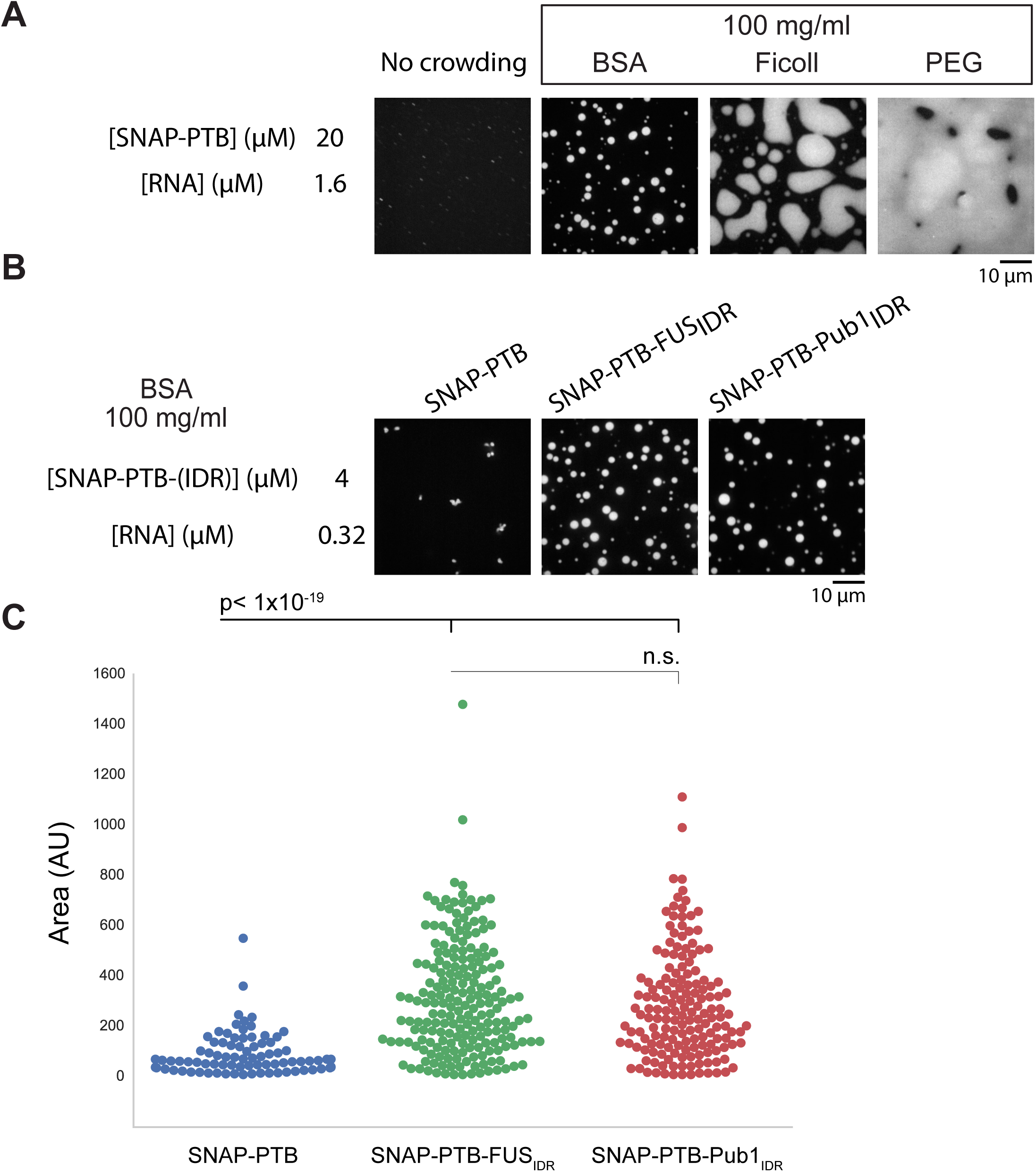
IDRs Enhance LLPS of PTB Plus RNA in the Presence of BSA. (A) Fluorescent microscopy images of phase separated droplets formed by SNAP-PTB and RNA in the presence or absence of 100 mg/ml BSA, Ficoll, or PEG. (B) Fluorescent microscopy images of 4 uM SNAP-PTB, SNAP-PTB-FUS_IDR_, or SNAP-PTB-Pub1_IDR_ plus 0.32 uM RNA assemblies in the presence or absence of 100 mg/ml BSA. (C) Quantification of assembly area for (B), with arbitrary units. Significance calculated with Welch’s *t*-test for unequal size and variance.

A second prediction of the model is that while IDRs alone are not sufficient to drive phase separation in the presence of competitor proteins, IDRs would contribute binding energy to phase separation driven by specific interactions, decreasing the threshold concentration of assembly. To test this prediction, we examined how IDRs affect PTB-RNA phase separation in the presence of competitor proteins. For example, 4 uM PTB and 0.32 uM RNA do not phase separate in 100 mg/ml BSA. However, we observed LLPS with identical concentrations of RNA and PTB, when the PTB was fused to either the FUS or Pub1 IDR **(Figure 4B)**. PTB fused to either IDR showed an increase in both the number and size of the assemblies visualized (**Figure 4C**). Therefore, weak interactions of IDRs can enhance phase separation in the presence of competitor proteins, when present in molecules that also contain specific interactions which are less susceptible to competition from cellular macromolecules.

### IDRs are often neither sufficient nor necessary *in vivo* to target components to RNP granules

An assembly mechanism for RNP granules driven by specific interactions aided by promiscuous interactions of IDRs has predictions for how components would be recruited to RNP granules. Specifically, one would predict that generally IDRs would not be sufficient to target a protein to an RNP granule, unless they contained a specific SLiM. Moreover, IDRs would not be required for recruitment to a granule, although they could affect the partition coefficient (the concentration of a component within versus outside of a granule).

To examine how IDRs of yeast proteins affect their targeting to P-bodies, we examined if IDRs within Lsm4, Dhh1, Pop2, and Ccr4 **(Figure 5A**) were necessary and/or sufficient for their recruitment into P-bodies. The IDRs of Lsm4, Dhh1, Pop2, and Ccr4 were fused separately to either GFP or mCherry. IDR-fusion proteins were expressed in yeast co-expressing a chromosomally GFP-tagged P-body component or containing a secondary plasmid containing a mCherry tagged P-body component. P-bodies were induced by glucose deprivation for 15 minutes, and the percentage of P-bodies containing the IDR fusion protein was counted. For example, clear enrichment in P-bodies was detectable for full length Lsm4 (**Figure 5B**). However, the Lsm4 IDR was not sufficient for P-body localization **(Figure 5B)**. Similarly, the IDRs of Dhh1, Pop2, and Ccr4, were insufficient for recruitment to P-bodies **(Figure 5C).** We then removed these IDRs from their full-length proteins, and found that deletion of the IDRs in Lsm4, Dhh1, Ccr4 had little to no effect on their recruitment to P-bodies **(Figure 5B,C)**. However, for the already poorly localized Pop2, deletion of the IDR did have a noticeable impact on localization (Figure 5C). Thus, the IDRs of Lsm4, Dhh1, Ccr4, and Pop2 are not sufficient their recruitment into P-bodies, but may contribute in cases where recruitment is already poor such as Pop2.

**Figure 5.**
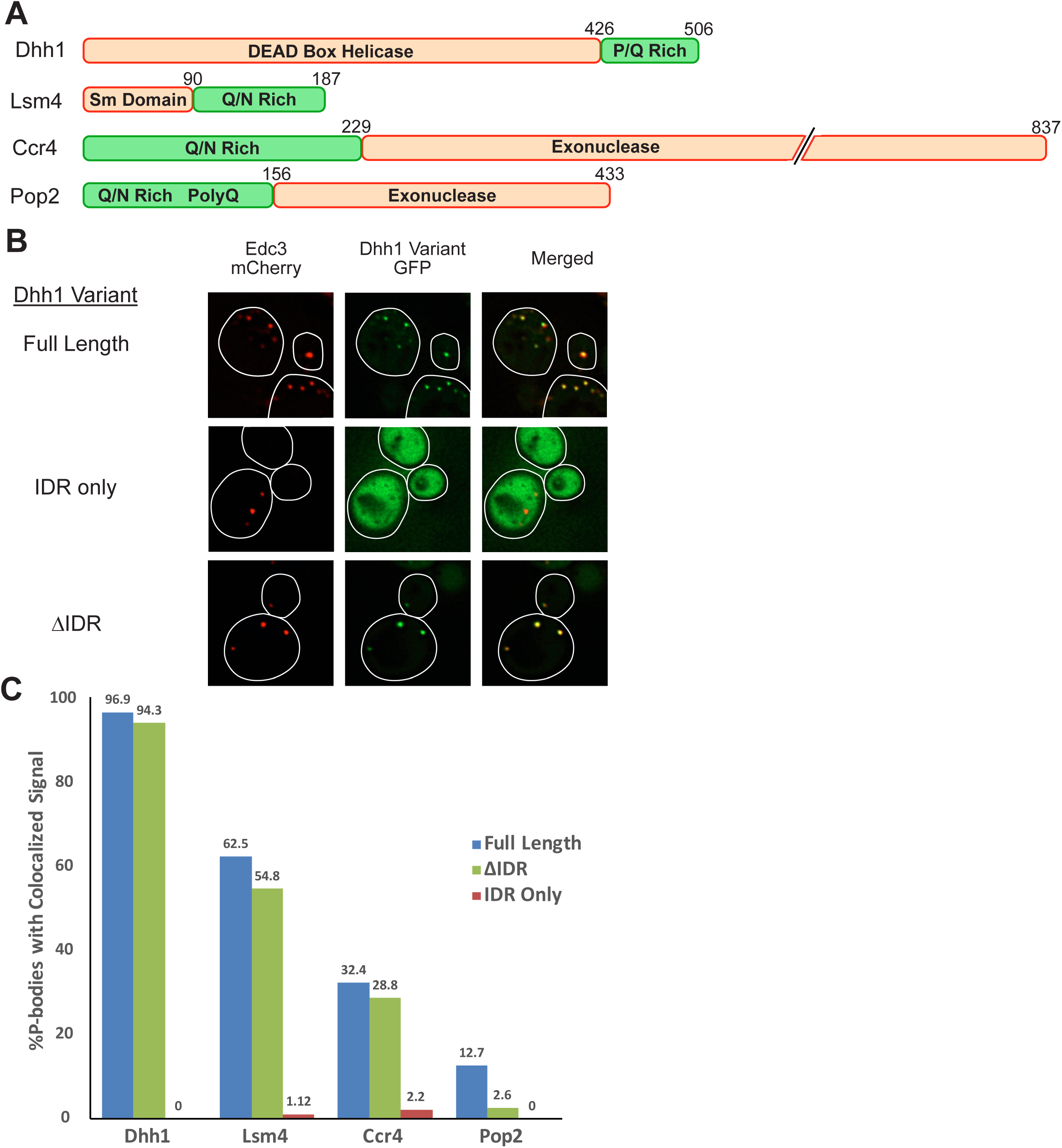
IDRs are neither sufficient nor required for P-body localization. (A) Domain structures of the yeast proteins Dhh1, Lsm4, Ccr4 and Pop2 (B) Dhh1-GFP variant fusions were expressed in Edc3-mCherry expressing yeast. After 10 minutes of glucose deprivation to induce P-bodies, cells were visualized by fluorescence microscopy. Representative images are presented. (C) Quantification of the percentage of P-bodies that exhibited colocalization with the expressed fusion protein. GFP was fused to the N-terminus of Dhh1, Ccr4, and Pop2 variants. These variants were cotransformed with Edc3-mCherry. mCherry was fused to the C-terminus of the Lsm4 variants, which were expressed in cells encoding genomically-tagged Dcp2-GFP. Dhh1ΔIDR 1-427, Dhh1 IDR 427-506; Ccr4ΔIDR 148-837, Ccr4 IDR 1-229; Pop2ΔIDR 147-433, Pop2 IDR 1-156; Lsm4ΔIDR 1-90, Lsm4 IDR 91-187.

### IDRs can enhance LLPS driven by specific interactions in cells

The observations above suggest cellular assemblies such as RNP granules may form with assembly primarily driven by a set of specific interactions, with the prevalence of IDR regions in such assemblies contributing either a second set of promiscuous nonspecific interactions that would enhance assembly, or having specific interactions with themselves that require high local concentrations to form. Reported observations suggest, however, that some IDRs noticeably contribute to RNP granule assembly in genetic backgrounds that limit assembly. One example of this phenomenon is the previous observation that the C-terminal IDR of Lsm4 is not required for P-body assembly normally, but plays a role in a strain lacking the P-body scaffold protein Edc3 (Decker et al., 2007).

To determine if this may be a more general phenomenon, we examined how the C-terminal IDR of the yeast Dhh1 protein promotes P-body formation. In *edc3*Δ *lsm4*Δ*C* yeast strains, which lack visible P bodies, P-body formation can be partially rescued by the addition of a single copy plasmid providing an extra copy of the Dhh1 gene, which through specific interactions with RNA and Pat1 enhances P-body assembly (Rao et al., 2017, submitted). Overexpression of Dhh1 in an *edc3*Δ *lsm4*Δ*C* background creates a cellular context where P-bodies are just above the threshold for assembly. Dhh1 also has a C-terminal P/Q rich IDR (**Figure 4**). To determine whether this C-terminal IDR contributes to P-body assembly, we compared the ability of full length Dhh1 and a Dhh1*Δ*IDR truncation (1-427), which lacks the C-terminal IDR (residues 428 to 506), to rescue P-body formation in an *edc3*Δ *lsm4*Δ*C* strain.

We found that wild-type Dhh1 rescues P-body formation in the *edc3*Δ *lsm4*Δ*C* strain, yet the Dhh1*Δ*IDR variant fails to do so (**Figure 6A,B**), despite being expressed at levels similar to the full-length protein **(Figure S2A)**. This demonstrates that the C-terminal IDR of Dhh1, while not required for P-body formation normally, can contribute additional interactions that enhance the formation of P-bodies when granule assembly is partially impaired.

**Figure 6.**
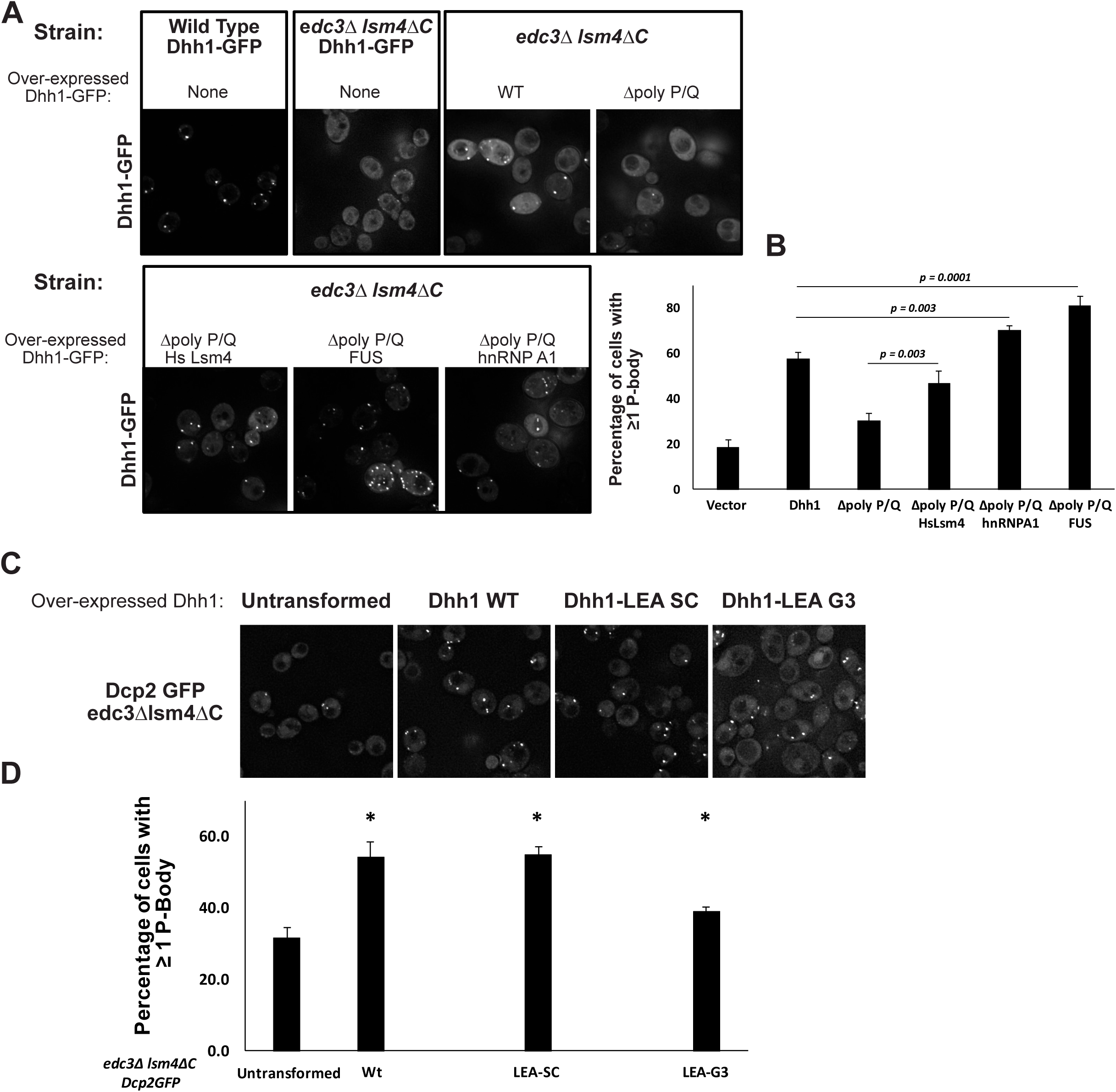
Specific interactions can synergize with promiscuous nonspecific interactions to drive assembly. (A) Fluorescent microscopy images of cells expressing Dhh1-GFP, either genomically or as a plasmid-expressed Dhh1-GFP variant. Cells were deprived of glucose for 10 minutes to induced P-body assembly. (B) Quantification of (A), depicting the percentage of cells containing at least one P-body. (Student’s *t*-test) (C) Fluorescent microscopy images of cells expressing Dhh1 – LEA variants or wild-type Dhh1. Cells were deprived of glucose for 10 minutes to induced P-body assembly, and visualized by genomically GFP-tagged Dcp2. (D) Quantification of (C), depicting the percentage of cells containing at least one P-body. (Student’s *t*-test, * p<0.05)

In principle, the Dhh1-IDR could provide a specific interaction, perhaps containing a SLiM, or a promiscuous interaction as we observed for several IDRs *in vitro*. If the Dhh1-IDR makes a specific interaction, then it should not be functionally replaceable by other IDRs capable of promiscuous interactions. Alternatively, if this IDR simply provides additional promiscuous interactions then any IDR capable of such interactions should functionally replace the Dhh1-IDR in promoting P-body assembly. To distinguish between these possibilities, we determined whether the IDRs of human Lsm4, a P-body component, as well as the IDRs of two human stress granule components, hnRNPA1, and the N-terminal domain of FUS, could replace the function of the Dhh1 IDR. We also tested the disordered regions of two Late-Embryogenesis Abundant (LEA) - like proteins, which are proposed to provide desiccation protection by interacting promiscuously with proteins in the cell, potentially in lieu of water (Hand et al., 2011). We utilized the IDRs of human proteins because these are very unlikely to contain specific binding partners in yeast.

All three granule-component IDRs complemented the P-body assembly defect seen in the Dhh1-ΔIDR construct (**Figure 6B**). Additionally, 2 LEA-like protein (LEA Group 3 – like from the brine shrimp *Artemia franciscana,* “LEA-G3,” and LEA Group2 – like from the nematode *Steinernema carpocapsae*, “LEA-SC”) IDRs also rescued the assembly defect (**Figure 6C**). Addition of these IDRs does not cause appreciable assembly of large structures without glucose deprivation (**Figure S2B**), demonstrating these assemblies are indeed P-bodies and not a different constitutive aggregate. These results argue that the Dhh1 IDR does not provide a specific interaction necessary for P-body assembly, as it can be replaced with a variety of other human IDRs. This result also demonstrates that multiple different IDRs can complement the Dhh1ΔIDR, consistent with promiscuous, nonspecific interactions of the IDRs contributing to RNP granule assembly in conjunction with specific interactions.

## DISCUSSION

RNP granules are cytoplasmic assemblies composed of specific groups of cellular proteins and RNA molecules (Jain et al., 2016, Khong et al., 2017 in submission). In principle, a specific assembly could be assembled in three manners: a) solely a set of specific interactions with well defined, and limited binding partners; b) through a summation of promiscuous interactions, where the sum of this network of interactions for a given molecule would bias its assembly characteristics, or c) through a combination of specific and promiscuous interactions. This third potential mechanism is supported by genetic analyses of the interactions that drive RNP granule assembly as well as our own findings. Specific interactions can clearly be important for assembly. For example, Edc3 dimerization via its YjeF-N domain is important for P-body assembly in yeast (Decker et al., 2007). G3BP dimerization, as well as interactions with caprin, are important for mammalian stress granule assembly (Tourriere et al., 2003; Kedersha et al., 2016). Some specific interactions can involve SLiMs found in IDRs that specifically interact with well-folded domains of other RNA binding proteins (reviewed in Jonas and Izaurralde, 2013). One example of this phenomena is the disruption of Edc3 localization to P-bodies in yeast caused by deletion of or interference with specific SLiMs in Dcp2’s C-terminal IDR, which interact with a surface of Edc3 (Fromm et al., 2012, 2014). Thus, specific interactions between RNA binding proteins play important roles in formation of P bodies and recruitment of molecules into them. However, we have also shown that promiscuous interactions can play a role in assembly.

A key contribution of this work is to provide evidence that at least some IDRs function to promote RNP granule assembly both in cells, and in model biochemical systems, through weak interactions that require being coupled to protein domains with specific interactions. First, examination of the FUS, hnRNPA1, and eIF4GII IDRs reveal that they all interact nonspecifically with generic proteins, and those proteins and yeast lysates disrupt their ability to undergo LLPS in isolation (**Figure 1 & 2**). However, when tethered to the PTB RNA binding protein, which phase separates in the presence of RNA, promiscuous IDRs can promote LLPS, even in the presence of competitor proteins (**Figure 4**). Third, the C-terminal P/Q rich IDR of Dhh1 promotes P-body assembly in yeast, and this domain can be replaced by the IDRs of human Lsm4, hnRNPA1, or FUS, or by specific LEA proteins from brine shrimp or nematodes (**Figure 6**). The contribution of such IDRs to assembly is likely due to the ability of IDRs to promote LLPS through a variety of weak promiscuous interactions including electrostatic, cation-π, dipole-dipole and π-π stacking interactions (Brangwynne et al., 2015; Nott et al., 2015), which would be enhanced through effects analogous to avidity by coupled specific interactions of adjacent domains (Jencks, 1981).

Additional evidence exists that LLPS can be driven by combined specific and non-specific interactions. For example, even very high expression levels of hnRNPA1-Cry2 or DDX4-Cry2 fusion proteins do not phase separate in cells, unless the Cry2 protein is first triggered to assemble through specific light activated interactions (See Figure 2B&C of Shin et al., 2016). This observation highlights how specific oligomerization domains can act cooperatively with IDRs to promote LLPS in cells, and how some IDRs may be insufficient to undergo LLPS without additional oligomerization elements. As an example of the importance of non-specific interactions in promoting cellular LLPS, the C-terminal IDR of yeast Lsm4 can enhance yeast P-body formation, but it can be functionally replaced in this role by other IDRs (Decker et al., 2007). Moreover, polyQ rich tracts, which are disordered IDRs capable of diverse interactions, are prevalent in P-body components and RNA binding proteins (Decker et al., 2007; Reijns et al., 2008), and can function in RNP granule assembly in *A. gossypii* (Lee et al., 2015). Taken together, we suggest that many IDRs on RNA binding proteins provide an additional layer of nonspecific interactions, and those interactions can contribute to granule formation when they synergize with more specific interactions to stabilize the macroscopic structure.

An important point is that, even when insufficient in themselves to promote LLPS, promiscuous IDRs can decrease the critical concentration for phase separation driven by more specific interactions. We demonstrate this phenomenon for phase separation of PTB and RNA *in vitro* (**Figure 4**), and for P-body assembly *in vivo* (**Figure 6**). This highlights that in a phase diagram describing an assembly based on specific and promiscuous interactions, the addition of promiscuous interactions can shift the system from an unassembled state to an assembled state (**Figure 7A,B**).

**Figure 7.**
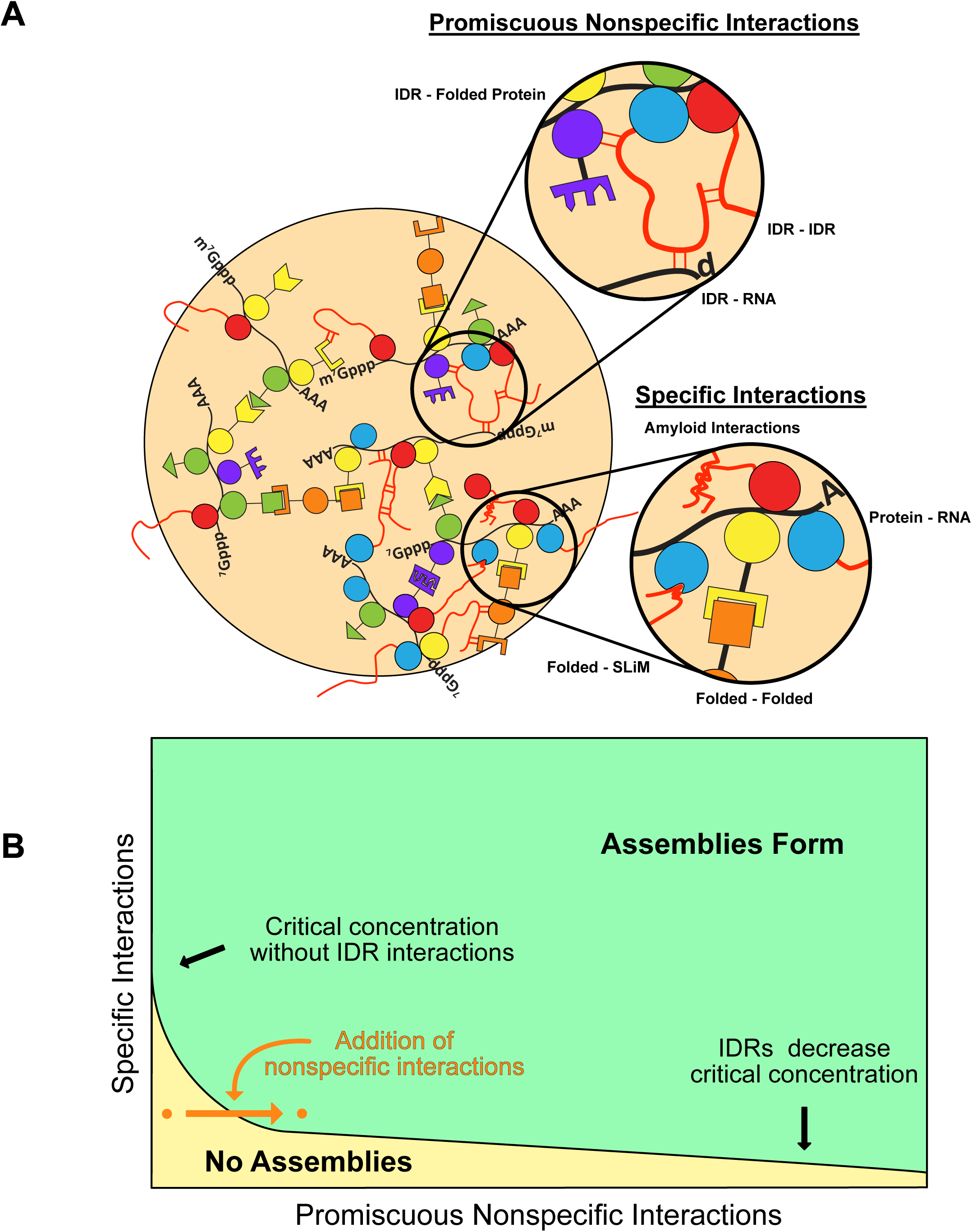
Model of RNP granule assembly and contributions of IDRs. (A) RNP granules assembly by a wide variety of specific and nonspecific interactions. (B) A theoretical phase diagram depicting how the addition of nonspecific, IDR-driven interactions could decrease the critical concentration of assembly for higher-order structures.

Not all IDRs will influence RNP granule formation in the same molecular manner. Some IDRs will provide specific interactions through SLiMs (Jonas and Izaurralde, 2013). Some IDRs may also afford interaction specificity through formation of local structure, including amyloid-like cross-beta interactions, that could be important in biological contexts where RNP granules need to be long-lived or mechanically stable (Boke and Mitchison, 2017; Kato et al., 2012) or *α*-helices (Conicella et al., 2016), both of which should show some sequence specificity. Charge patterning can also afford sequence specificity, although likely to a lower degree (Nott et al., 2015; Pak et al., 2016). Finally, as suggested here, some IDRs will provide promiscuous interactions that can enhance RNP granule assembly. Therefore, interactions undergone by any individual IDR that can contribute to intracellular LLPS likely lay on a scale from low affinity and highly promiscuous, to moderate affinity and selective.

*A priori*, there are three general classes of promiscuous interactions that IDRs could contribute to granule assembly. First, IDRs could interact with themselves or with other IDRs through weak interactions, which is suggested by observations that both IDR-based hydrogels and phase separated liquid droplets can recruit proteins with different IDRs (Kato et al., 2012; Lin et al., 2015). Second, IDRs could have promiscuous interactions with RNAs, which is suggested by observations that some IDRs cross-link to RNA *in vivo* (Castello et al., 2016) and some IDRs bind RNA *in vitro* (Lin et al., 2015; Molliex et al., 2015). Finally, IDRs could make promiscuous interactions with other well-folded domains of granule components. Note that promiscuous interactions of IDRs with well-folded domains of proteins could provide an evolutionary starting point for the formation of SLiMs, which are often found in IDRs. An important future goal will be in determining how IDRs utilize each of these interaction types to contribution to granule formation.

Utilizing promiscuous nonspecific interactions of IDRs to modulate the assembly of macro-scale complexes has unique advantages. First, since such nonspecific interactions are not limited to defined components or stereospecific arrangements, they can interact promiscuously with any number of individual components to enhance assembly. For example, in RNP granules, a diversity of mRNPs with different RNA binding proteins can be components of the granule. Promiscuous IDRs on RNA-binding proteins could interact with any of these mRNPs to enhance granule assembly. Moreover, IDRs can be subject to rapid evolution, and control by post-translational modifications, thus making them ideal components to change granule assembly parameters under selective pressure and in response to signaling pathways. Finally, we note that because higher-order assemblies are large with respect to a single IDR, promiscuous interactions of IDRs will mostly occur within the quinary space of the assembly, rather than with proteins outside of the assembly. This makes large assemblies particularly well suited to enhancement by IDRs.

Macromolecular assembly and concomitant LLPS mediated by combinations of specific and promiscuous interactions is a general mechanism for forming dynamic, meso-scale structures in eukaryotic cells. Eukaryotic cells contain many such assemblies including RNP granules, signaling complexes, DNA damage repair foci, and transcription complexes. It is notable that components of all of these assemblies are enriched in IDRs (Banani et al., 2016; Hegde et al., 2010; Hnisz et al., 2017; Iakoucheva et al., 2002; Kai, 2016; Minezaki et al., 2006). Thus, we suggest that higher order complexes will often be assembled by a combination of specific interactions that drive assembly, reinforced by a network of promiscuous nonspecific IDR based interactions, which stabilize the complex because of their physical coupling to specific assembly components. Such assemblies will be easily modified over time via evolution, or in a dynamic sense by signaling pathways and post-translational modification. This would occur without having to change the underlying specific assembly interactions, thus allowing both rapid evolution of and immediate control over intracellular assemblies.

### Author Contributions

D.S.W.P. designed and performed experiments, provided substantial intellectual input, contributed to writing the paper, and edited the paper.

B.V.T. and B.R. designed and performed experiments, and edited the paper.

Y.L. designed and performed experiments, provided substantial intellectual input, and edited the paper.

L.M. performed experiments.

M.K.R. Provided substantial intellectual input, assisted with experimental design, and edited the paper.

R.P. provided substantial intellectual input, assisted with experimental design, contributed to writing the paper, and edited the paper.

## EXPERIMENTAL PROCEDURES

### Protein Purification and Labeling

Proteins were expressed and purified as previously reported (Lin et al., 2015). Proteins were expressed from the pMal-c2 vector (NEB), except for full length hnRNPA1 and related mutants, which were cloned into a modified pet11a vector (Novagen). Proteins were expressed in *E. coli* BL21(DE3) and purified with Ni-NTA and/or amylose resin under standard conditions. SNAP-PTB-IDRs were further purified through a Superdex200 column (GE Healthcare). Proteins were fluorescently-labeled with SNAP-Surface 488 or SNAP-Surface 649 (NEB) according to the manufacturer’s protocols. Unincorporated dye was removed using Zeba Spin Desalting Columns, 7K MWCO (Thermo Fisher). Proteins were concentrated using Amicon Ultra 10K MWCO centrifugal filters (Milipore) and aggregates removed by ultra-centrifugation at 4°C for 30’ at 50K RPM in a Beckman-Coulter TLA 100.2 rotor.

### Fluorescence Microscopy

All yeast experiments and all images of SNAP-IDR and SNAP-hnRNPA1 were acquired on a DeltaVision epi-fluorescence microscope, equipped with an SCMOS camera. All images of SNAP-PTB-IDR were acquired on a Leica-based spinning disk confocal microscope (EMCCD digital camera, ImagEM X2, Hamamatsu; confocal scanner unit, CSU-X1, Yokogawa).

### Droplet Assembly

For SNAP-IDRs and SNAP-hnRNPA1 (~2% fluorescently labeled), droplet assembly was initiated by diluting solutions to 37.5 mM NaCl, 20 mM Tris pH 7.4, 1 mM DTT. For SNAP-PTB-IDRs, proteins and RNA, (UCUCUAAAAA)_5_, were mixed at the indicated concentrations (including 100 nM SNAP-PTB-IDRs labeled with SNAP-Surface 649) in 100 mM NaCl, 20 mM imidazole pH 7.0, 1 mM DTT, 10% glycerol. N-terminal purification tags of SNAP-hnRNPA1 were removed by HRV C3 protease (EMD Milipore) during the dye conjugation step (Lin et al., 2015). N-terminal MBP and C-terminal His tags of SNAP-IDRs and SNAP-PTB-IDRs were cleaved just prior to droplet assembly with TEV protease (Promega ProTEV). Reactions were performed in glass-bottom chambers passivated with 3% BSA. FITC-conjugated Lysozyme (Nanocs) and FITC-BSA (Thermo Fisher Scientific) were mixed with SNAP fusion proteins prior to droplet assembly at concentrations of 100 nM and 10 nM, respectively.

### Droplet Quantification

Images were analyzed in FIJI as follows. Images were imported, flattened with a maximal intensity projection when applicable (hnRNP A1Δhexa droplets), and then thresholded using either the Default method (hnRNP A1 Δhexa droplets) or the Otsu method (PTB droplets) (‘Threshold’). Binary images were eroded (‘Erode) once to remove single pixels, then dialated (‘Dialate’) once to return droplets to their original size. This was followed by watershedding (‘Watershed’) to separate proximal droplets. FIJIs ‘Analyze Particles’ was used to generate ROIs, which were used to measure the maximal intensity projection, generating area and mean intensity values for each assembly.

### Microscopy and Quantification for P-body Colocalization

Cells were grown at 30°C to OD_600_ of 0.3-0.5 in minimal media with 2% glucose as a carbon source and with necessary amino acid dropout to maintain plasmids and express constructs (See Supplemental File “Strains Plasmids and Antibodies”). Cells were stressed by glucose deprivation for 15 minutes before cells were concentrated for immediate microscopic examination at room temperature. All images underwent deconvolution using DeltaVision’s algorithm.

Images were quantified using FIJI. To optimize yeast colocalization accuracy, single plane images were used and analysis were done in a blind manner. P-bodies were identified using protein markers (either Dcp2-GFP, Edc3-mcherry). Corresponding enrichment of the construct within the P bodies was then assessed manually. Manual assessment was required due to differential strengths of cytoplasmic signals between cells arising from stochastic variation and/or potentially different copy numbers of plasmids between cells.

### Growth and Microscopy of Dhh1 Variants

To test the effect of Dhh1-IDR chimera on P-body recovery in the edc3Δ lsm4ΔC yeast (Strain yRP2338), yeast were transformed with vector only or vectors containing GFP fusions of Dhh1 wt, Dhh1-1-427 and Dhh1-IDR chimera using standard yeast transformation protocols. Two individual transformants were selected as biological replicates. The replicates were grown overnight to saturation at 30 ^o^C with shaking in SD-Ura media (minimal media), containing 2% dextrose. The saturated cultures were re-inoculated into fresh SD-Ura media and grown to OD = 0.4-0.5. The cells were pelleted and transferred to S-Ura media lacking dextrose and shaken at 30 ^o^C for 10 min prior to microscopic analysis. For the unstressed conditions, the cells were pelleted without glucose starvation.

Yeast were analyzed via fluorescence microscopy on the DeltaVision Elite microscope with a 100 X objective using a PCO Edge sCMOS camera. ≥2 images comprising of 9 Z-sections were obtained for each replicate. Images were analyzed using Image J. Z-projections derived from summation of the Z-sections with constant thresholding were used to count the number of yeast cells with ≥1 GFP-positive granule, and the percentage of cells with at least 1 granule was calculated.

### Plasmid construction

The Dhh1-GFP gene fragment containing the Dhh1 promoter was PCR amplified using the genomic DNA from the Dhh1-GFP yeast strain (yeast GFP collection) and BSR_DhhGFP416NF and BSR_DhhGFP416NR primers. The Adh1 terminator fragment was clone using the primers BSR_Adh1SacF and BSR_Adh1SacR. The Dhh1-GFP and Adh1 terminator fragments were inserted sequentially into the XhoI and SacI digested pRS416 vector, respectively, via Infusion cloning (Takara). The poly P/Q residues of Dhh1 (428-506 were deleted from the Dhh1-GFP containing vector using primers, Dhh11-427F and Dhh11-427R via the Phusion mutagenesis protocol (Thermo Fisher). Lastly, the intron-less IDR sequence for HsLsm4 was synthesized using gBLOCK technology from IDT technologies. The IDRs were PCR amplified using primers, BSR_427FUSF and BSR_427FUSR, BSR_427A1F and BSR_427A1R, BSR_427HsLsm4F and BSR_427HsLsm4R, for FUS, hnRNPA1 and HsLsm4, respectively and cloned into the linearized Dhh1-1-427-GFP vector using Infusion cloning.

---

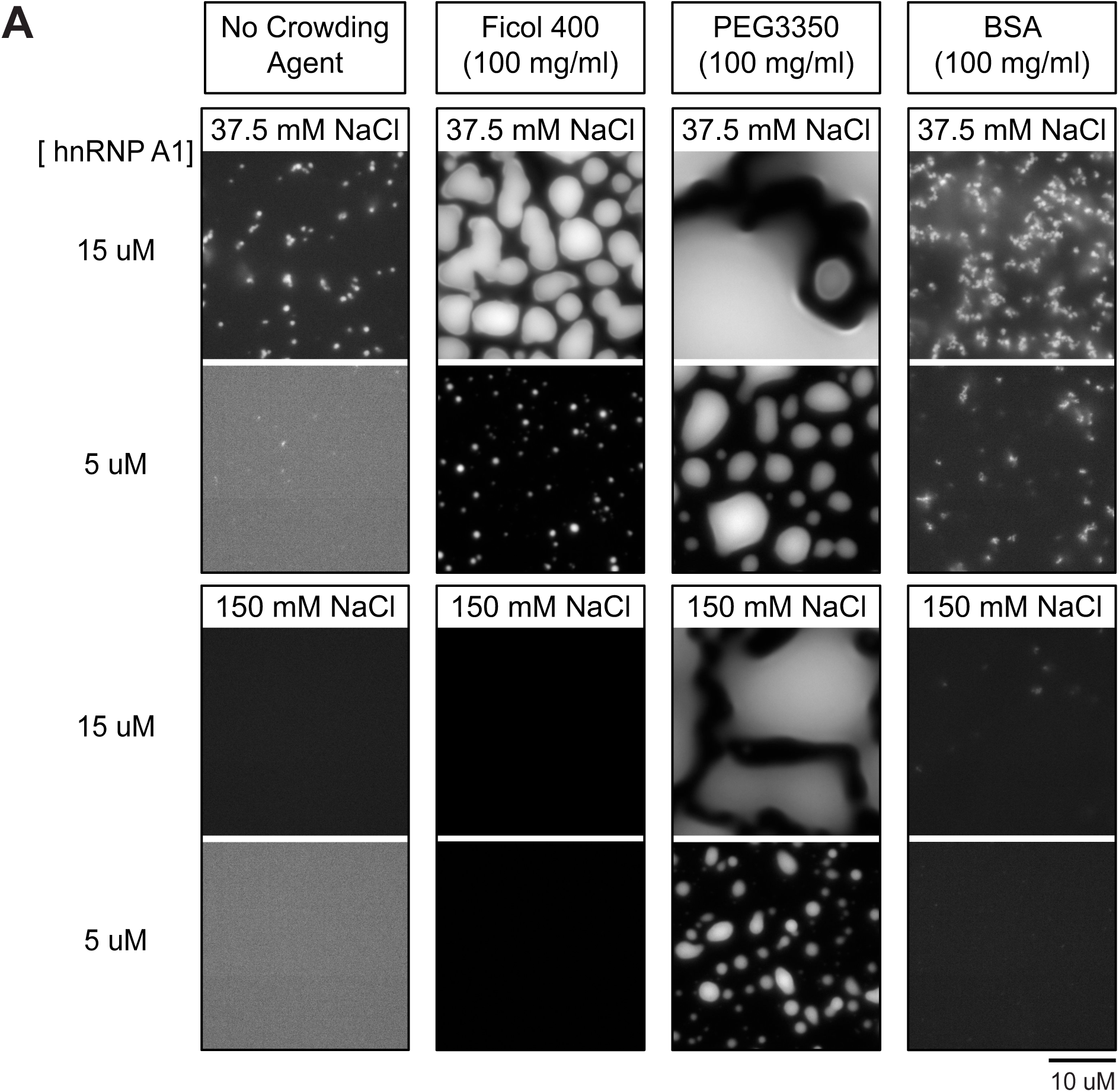
S1 Related to Figures 1 and 2: Diverse effects of crowding agents and proteins on LLPS. (A) Fluorescence microscope images of structures formed by SNAP-hnRNP A1 at either 150 mM NaCl or 37.5 mM NaCl in the absence or presence of 100 mg/ml Ficol 400, PEG 3350, or BSA. SNAP-hnRNP A1 concentrations were either 15 uM or 5 uM.

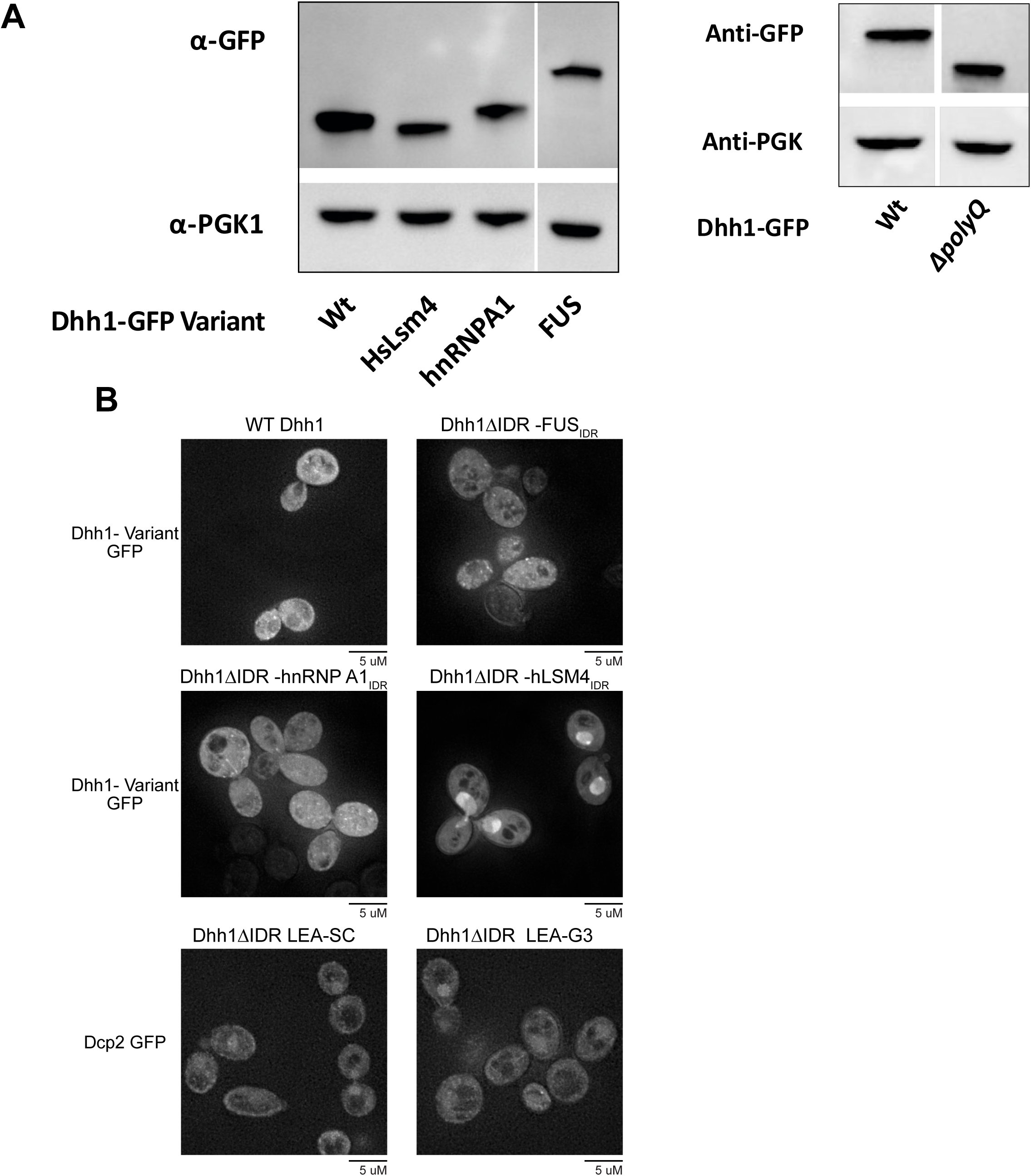
Sup 2 Related to Figure 6: Dhh1 IDR fusions do not form large assemblies in the absence of stress. (A) Western blot of Dhh1 variant expression (B) Fluorescent microscopy images of cells expressing Dhh1-GFP, either genomically or as a plasmid-expressed Dhh1-GFP variant. Cells were growing under log phase growth conditions just prior to imaging.

